# High-Throughput Screening of the *Saccharomyces cerevisiae* Genome for 2-Amino-3-Methylimidazo [4,5-*f*] Quinoline Resistance Identifies Colon Cancer-Associated Genes

**DOI:** 10.1101/2022.10.26.513854

**Authors:** Michael Dolan, Nick St. John, Faizan Zaidi, Francis Doyle, Michael Fasullo

**Affiliations:** Colleges of Nanoscale Science and Engineering, State University of New York Polytechnic Institute, 257 Fuller Drive, Albany, New York 12203

**Keywords:** genome profiling, budding yeast, heterocyclic aromatic amine, colorectal cancer

## Abstract

Heterocyclic aromatic amines (HAAs) are potent carcinogenic agents found in charred meats and cigarette smoke. However, few eukaryotic resistance genes have been identified. We used *Saccharomyces cerevisiae* (budding yeast) to identify genes that confer resistance to 2-amino-3-methylimidazo[4,5-f]quinoline (IQ). CYP1A2 and NAT2 activate IQ to become a mutagenic nitrenium compound. We introduced an expression vector that contains human CYP1A2 and NAT2 genes into selected mutant strains and the diploid yeast deletion collection. The deletion libraries expressing CYP1A2 and NAT2 or no human genes were exposed to either 400 or 800 μM IQ for five or ten generations. DNA barcodes were sequenced using the Illumina HiSeq 2500 platform and statistical significance was determined for exactly matched barcodes. Four screens for IQ resistance in the “humanized” collection identified 1160 unique ORFs, of which 337 were validated or duplicated in at least two screens. Two screens of the original yeast library identified 101 genes that overlap with the 337 previously identified. Selected genes were validated by growth curves, competitive growth assays, or trypan blue assays. Prominent among both sets are ribosomal protein genes, while nitrogen metabolism, cell wall synthesis, and phosphatase genes were identified among the “humanized” library. Protein complexes identified included the casein kinase 2 (CK2) and histone chaperone (HIR) complex. DNA repair genes included *NTG1, RAD18, RAD9, PSY2* and *UBC13*. Polymorphisms in human *NTHL1*, the *NTG1* ortholog, and *RAD18* are risk factors for colon cancer. These studies thus provoke questions of whether genetic risk factors for colon cancer confer more HAA-associated toxicity.

## INTRODUCTION

The heterocyclic aromatic amine (HAA) 2-amino-3-methylimidazo[4,5-*f*] quinoline (IQ) is a mutagen (Kasai *et al*., 1980) and a class 2A carcinogen (IARC, 1992; Adeyeye and Ashaolu, 2021), present in charred meat and cigarette smoke. IQ and other HAAs are generated by Maillard reactions involving creatine, sugar, and an amino acid (Jägerstad *et al*., 1983); these reactions occur when charring meat and suggest why red meat consumption increases colon cancer risk (Sinha *et al*., 1999; Sinha *et al*., 2001; Liniesen *et al*., 2002; Aykan, 2015; Martinez Gongora *et al*., 2019). In rats, IQ exposure induces liver, kidney, and colorectal cancers (Turesky *et al*., 1995), while in nonhuman primates, IQ exposure induces liver cancer. (Adamson *et al*., 1990; Turesky *et al*., 1996a). IQ exposure is associated with oxidative stress (Carvalho *et al*., 2015) and higher levels of DNA damage (Carvalho *et al*., 2016). Genotoxic endpoints of IQ exposure include higher frequencies of both mutation (Aeschbacher and Turesky, 1992) and sister chromatid exchange (Sawada *et al*., 1994). These studies suggest that IQ acts as a genotoxin contributing to the carcinogenicity associated with red meat or cigarette smoke in humans.

IQ pre-incubated with rat liver S9 is more mutagenic in the Ames assay than IQ alone (Kasai *et al*. 1980), indicating that IQ’s genotoxicity requires bioactivation. In humans, bioactivation occurs by a two-step mechanism (Figure 1; Kim and Guengerich, 2005). First, cytochrome P450 enzymes (CYPs), such as CYP1A2, hydroxylate IQ to form N-hydroxy-IQ, and second, N-acetyltransferases (NATs), such as NAT1 and NAT2, acetylate the N-hydroxy-IQ to form N-acetoxy-IQ (Figure 1). This latter compound is unstable and undergoes spontaneous heterolytic cleavage to generate a highly reactive nitrenium ion (Kato, 1986; Turesky and Vouros, 2004; Kim and Guengerich, 2005), which forms C^8^ and N^2^ Gua adducts. Interestingly, the N^2^ Gua adduct appears refractory to nucleotide excision repair (NER) and is bypassed by error-prone polymerase (Stavros *et al*., 2014), while the C^8^ adduct is more efficiently removed (Turesky *et al*., 1996b). NAT2 expression increases the mutagenicity of IQ in both CYP1A2-expressing strains of *Salmonella typhimurium* and in Chinese hamster ovary cells (CHO) (Grant *et al*., 1992; Josephy *et al*., 1995) and the recombinogenicity of IQ in CYP1A2-expressing strains of *Saccharomyces cerevisiae* (budding yeast, Paladino, 1999). These studies indicate that expression of both cytochrome P450 and NAT enzymes are required for maximum levels of IQ genotoxicity.

**Figure 1.**
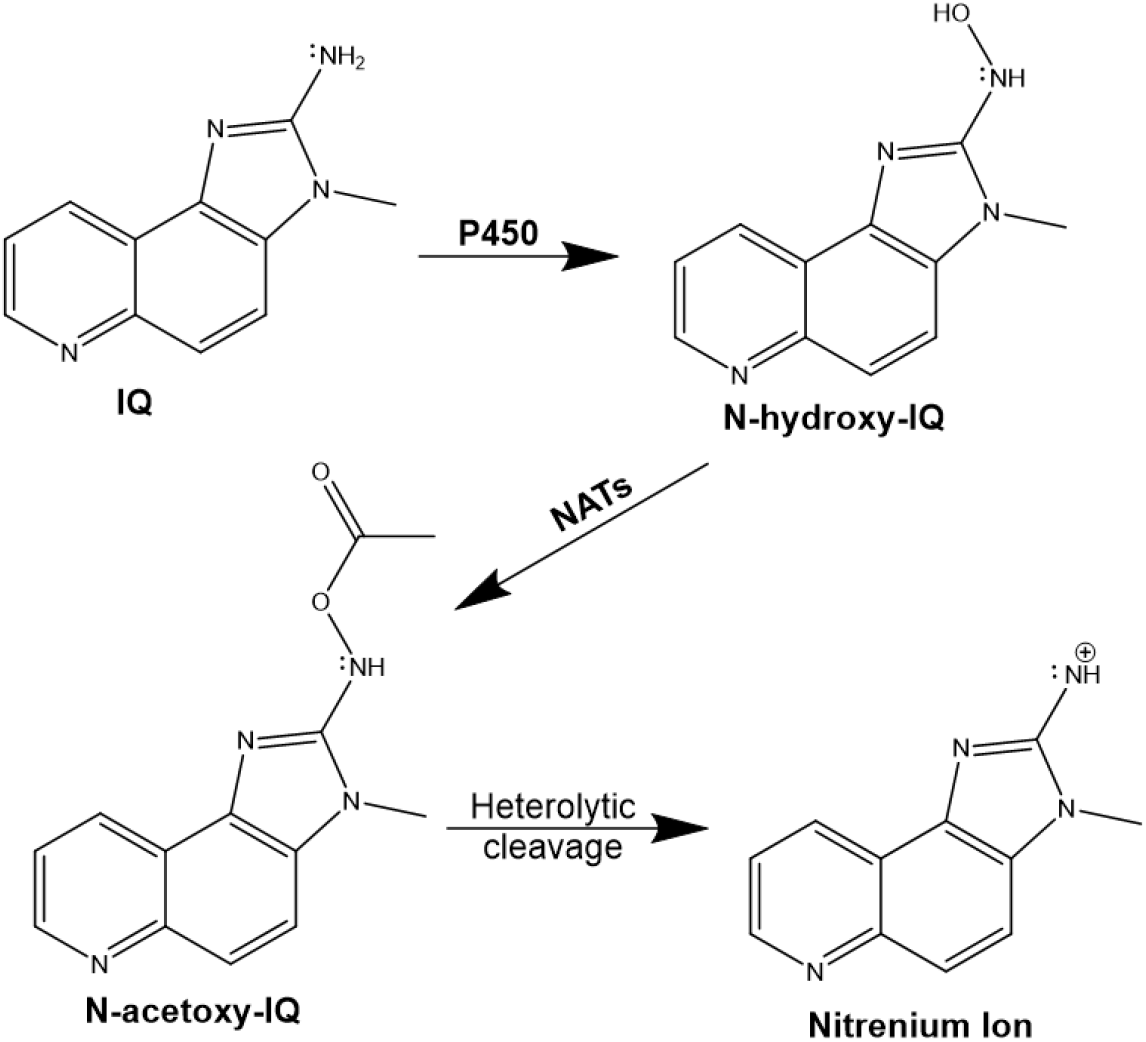
Bioactivation of the heterocyclic aromatic amine IQ by human CYP and NAT genes. The chemical structures of IQ, N-hydroxy-IQ, and N-acetoxy-IQ are shown in the figure. IQ is first hydroxylated by cytochrome P450 enzymes (CYPs) to form N-hydroxy-IQ. The hydroxylated IQ is then acetylated by N-acetyl transferases (NATs) to form N-acetoxy-IQ. This compound is unstable and generates the nitrenium ion shown in the figure.

Genetic factors that increase the mutagenicity of HAAs have been correlated to higher incidence of smoking and diet-associated cancers. For example, the human *NAT2*4* rapid acetylator allele, whose expression in CHO cells increases HAA-associated mutations more than that of *NAT2* slow acetylator alleles (Probst-Hensch *et al*., 1995; Grant *et al*., 1997; Fretland *et al*., 2001), is a colon cancer risk factor among Caucasians (Butler *et al*., 2008) and smokers (Nöthlings *et al*., 2009; Slattery *et al*., 1998). Individuals who frequently consume red meat and exhibit rapid acetylator NAT phenotypes, carrying *NAT2*4* or *NAT1*10* alleles, are at a higher risk for developing colon cancer (Metry *et al*., 2010; Lilla *et al*., 2006; Wang *et al*., 2011; Le Marchand, 2021). Although IQ is found in varying concentrations in the diet and cigarette smoke, the epidemiological studies suggest that IQ genotoxicity aggravates cancer risk.

Besides DNA adducts, IQ exposure also generates DNA strand breaks and oxidative stress, suggesting that DNA repair genes may function in suppressing IQ-associated genotoxicity (Pfau *et al*., 1999; Pezdirc *et al*., 2013). The importance of this question is underscored by observations that high risk factors for colon cancer include biallelic mutations of two base excision repair genes, *NTHL1* (Weren *et al*., 2015; Te Paske 2020) and *hMYH* (Wooden *et al*., 2004). Low penetrant colon cancer risk factors include polymorphisms in the DNA repair genes *RAD18, ERCC5, XPC, PARP*, and *APE1* (Pan, et al., 2012; Aggarwal *et al*., 2017; Matejcic *et al*., 2021). Whether DNA repair gene polymorphisms confer higher levels of HAA genotoxicity is unknown.

Budding yeast is a model organism for profiling genes that confer resistance to genotoxins. Over 30% of the yeast genome is orthologous to humans (O’Brien *et al*., 2005), and human orthologs can functionally substitute for 469 essential yeast genes (Kachroo *et al*., 2015). Each nonessential gene has been individually knocked out and tagged with both upsteam and downstream barcodes (uptags and downtags, Giaever *et al*., 2002); this deletion library has already been shown to be a powerful reagent in identifying drug targets and toxicant resistance genes (Jo *et al*., 2009; Giaever and Nislow, 2014; De La Rosa *et al*., 2017). While budding yeast does not bioactivate HAAs, expression of human CYP1A2 and NAT2 increases the HAA-associated translocation frequencies in budding yeast (Paladino *et al*., 1999). A “humanized” yeast deletion library expressing CYP1A2 and NAT2 has already been used to profile the yeast genome for aflatoxin B_1_ (AFB_1_) resistance (St. John *et al*., 2020). In this study, we profiled the same “humanized” deletion collection for resistance to IQ. We chose IQ because, among HAAs, activated IQ is the most recombinogenic in yeast (Paladino *et al*., 1999) and is highly mutagenic in strains of *S. typhimurium* (Knasmüller *et al*., 1999; Nohmi and Watanabe, 2021). We identified budding yeast genes that confer IQ resistance; mammalian orthologues of a subset of the IQ resistance genes are associated risk factors for colon cancer.

## MATERIALS AND METHODS

### Media and chemicals

Standard media were used for the culture of yeast and bacterial strains (Burke *et al*.,2000). The bacterial DH1 strains containing the vectors pCS316 and pCYP1A2_NAT2 were cultured in LB-Amp (Luria Broth media with 100 μg/mL ampicillin). Media used for the culture of yeast cells included YPD (yeast extract, peptone, dextrose), SC (synthetic complete, dextrose), and SC-URA (SC lacking uracil). Media to select for 5-fluoroorotic acid (FOA) resistance contained SC-URA supplemented with 4x uracil and FOA (750 μg/ml).

A monoclonal antibody to CYP1A2 (ab22717) and a polyclonal antibody to NAT2 (ab88443) were purchased from Abcam. Purified NAT2 protein was purchased from OriGene (CAT#: TP761755). 2-amino-3-methylimidazo [4,5-*f*] quinoline (IQ) was purchased from Toronto Research Chemicals. Methanol was purchased from Sigma. IQ was dissolved in methanol containing 0.1% acetic acid (MeOH).

### Strains and plasmids

Yeast strains used in this study were derived from either BY4741, BY4743 (Brachmann *et al*., 1998), or YA289 (St. John *et al*., 2020) and are of the S288C background. Strain genotypes are listed in Table S1. The yeast deletion collection is derived from BY4743, whose genotype is *MAT**a**/α his3Δ1/his3Δ1 leu2Δ0/leu2Δ0 LYS2/lys2Δ0 met15Δ0/MET15 ura3Δ0/ura3Δ0*. The diploid and haploid homozygous deletion libraries were purchased from Open Biosystems and are now available from Horizon Discovery (https://horizondiscovery.com/en/non-mammalian-research-tools/products/yeast-knockout). The pooled diploid homozygous deletion library (n = 4607) was a gift from C. Vulpe (University of Florida). The CYP1A2 expression plasmid, pCS316 (Sengstag *et al*., 1996), was obtained by CsCl centrifugation (Ausubel *et al*., 1995). The plasmid used in this study, pCYP1A2_NAT2 (St. John *et al*., 2020), was constructed by removing the hOR sequence from pCS316 and replacing it with the Not1 fragment containing human NAT2 from pRS424-CYP1A2_NAT2. The cDNA of NAT2 was previously positioned in the pGAC.24 (Lesser and Guthrie, 1994) expression plasmid under the control of the glycerol phosphate dehydrogenase (GPD) promoter.

### Transformation protocol

The modified diploid homozygous deletion libraries were prepared as previously described (St. John *et al*., 2020). In brief, FOA^R^ isolates were taken from each strain in the diploid homozygous deletion collection and inoculated in 100 μL of YPD in 96-well plates. After overnight incubation at 30 °C, plates were centrifuged, washed, and resuspended in one-step buffer (50% PEG, MW 3300, 0.2 N lithium acetate, 100 mM dithiothreitol, 500 μg/mL denatured salmon sperm DNA). Each well received 1 μg of pCYP1A2_NAT2 and plates were incubated for 30 mins at 30 °C, before pipetting 10 μL onto duplicate SC-URA plates. Two Ura^+^ transformants were chosen from each strain and frozen in SC-URA with 7.5% DMSO in 96-well plates. This strategy was successful in transforming approximately 90% of the diploid homozygous deletion library.

### Functional profiling of the yeast genome

Tubes of the pooled library (10 plates per tube) were thawed on ice and 250 μL of each tube’s content were added to 4.5 mL of YPD or SC-URA in a 15 mL tube and allowed to recover by incubation at 30 °C for two hours in a shaking incubator. From the recovered pooled libraries, 100 μL was inoculated into culture tubes containing 2 mL of either YPD for the original yeast library or SC-URA for the yeast deletion collection containing the pCYP1A2_NAT2 plasmid. Samples from each library were then treated in quadruplicate with 2% MeOH, 400 μM IQ, or 800 μM IQ. Samples were incubated on a rotary incubator at 30 °C for either 15 hours (for the YPD cultures) or 10 hours or 20 hours (for the SC-URA cultures), approximately five (5G) and 10 generations (10G). The cells were then centrifuged and washed thrice with sterile water.

To amplify molecular barcodes, DNA from each treatment tube was isolated using the “smash and grab” method (Hoffman and Winston, 1987) and resuspended in TE buffer (10 mM Tris-HCl, 1 mM EDTA, pH 8.0) at concentrations of 2-4 μg/μL. Polymerase chain reaction (PCR) was performed on each sample using customized oligos (Table S2) that hybridize to the regions just upstream and downstream of the barcode “uptag” sequences; the oligos contained a multiplex index and sequencing primers for use with the Illumina Hi-Seq 2500 (Robinson *et al*.,2014; Smith *et al*., 2009). PCR protocols for barcode amplification were previously described (Robinson, *et al*., 2014). The ~150 bp PCR products were resolved on a 10% polyacrylamide nondenaturing gel, extracted using the “crush and soak” method (Green and Sambrook, 2019), and quantified. A pooled sample containing 50 ng of each of the 32 samples was sent to the Bauer Sequencing Core Facility at Harvard University for quality assurance and sequencing using the Illumina HiSeq 2500 platform. For experiments comparing 5-generation vs 10-generation IQ exposures, samples were sent to the University at Buffalo Genomics and Bioinformatics Core Facility for quality assurance and sequencing on the Illumina HiSeq 2000 platform.

### Barcode analysis

The sequencing data obtained in the FASTQ files was demultiplexed to separate reads from different exposures. The individual files were then subjected to a FastQC report, then trimmed to include only the barcode region with 2-3 extra bases on either side. Barcodes were counted with an in-house program and associated with strains. Replicate treatments were grouped and compared to solvent treatment. Comparisons of treatment were performed using Bioconductor’s “TCC” package in RStudio (Sun *et al*., 2013), processed in batches using RStudio version 1.4.1717 and R version 4.1.0 using the code in File S1.

To determine which strains decreased in abundance after IQ exposure compared to MeOH exposure, m-values (log_2_ [barcode counts IQ-treated/barcode counts MeOH-treated]) were calculated. Negative m-values indicate that the strain was depleted after exposure and the corresponding ORF or gene conferred IQ resistance. Positive m-values indicate that the strain was enriched after treatment and the corresponding ORF conferred IQ sensitivity. Q-values (corrected p-values, Sun *et al*., 2013) less than 0.1 were deemed statistically significant.

### Western Blots

Expression of CYP1A2 and NAT2 in yeast cells was determined by Western blot. Cells were inoculated in YPD or SC-URA media, grown to log phase (A_600_ = 0.5-1), and then concentrated. Protein extracts were prepared as described previously by Foiani *et al*. (1994). Proteins were then separated on a 10% polyacrylamide gel and transferred to polyvinylidene fluoride membranes. To detect human CYP1A2, a mouse anti-CYP1A2 antibody (1:1,000) was used, followed by a goat anti-mouse secondary antibody (1:10,000). To detect human NAT2, a polyclonal mouse anti-NAT2 antibody (1:1,000) was used, followed by the same goat anti-mouse secondary antibody (1:10,000). Precision Plus Protein Western C Blotting Standards (Bio-Rad) were used for molecular weight standards.

### CYP1A2 and NAT2 enzyme activity assays

To determine whether the expressed protein was functional, the methoxyresorufin demethylase (MROD) activity of CYP1A2 was measured, using a method previously described (Fasullo *et al*., 2014). Microsomes were prepared using a modified protocol originally described by Pompon *et al*. (1996) and St. John *et al*. (2020). Microsomal protein concentrations were 15-20 mg/ml. The reaction mix consisted of adding 1 μL microsomes to 5 μM 7-methoxyresorufin in 10 mM Tris, pH 7.4 at 37 °C. Reactions were started with the addition of 500 μM NADPH and fluorescence was recorded every minute for one hour. Fluorescence of resorufin was measured on a Tecan Infinite M200 plate reader with an excitation wavelength of 535 nm and emission wavelength of 580 nm. Microsomes derived from BY4743 containing no CYP1A2 were used for the negative control. For a positive control, rat liver S9 fractions were used.

To measure the N-acetyl transferase activity of NAT2, we used cytosolic extracts from yeast expressing CYP1A2 and NAT2, prepared as described by Paladino *et al*. (1999). Briefly, a 100 mL culture, grown to mid-log phase, was pelleted and cells washed twice before being resuspended in 2 mL ice-cold disruption buffer (100 mM Tris, pH 7.4, 100 mM DTT, 1 mM EDTA, 10% (v/v) glycerol, and 1x yeast proteinase inhibitor cocktail IV (Thermo Scientific). Cells were lysed in the bullet blender after addition of 200 mg 0.45-0.55 mm glass beads. After 10 mins on the bullet blender, lysate was removed to a new tube and the beads were washed twice in an equal volume of ice-cold disruption buffer. The three lysate fractions were pooled in a new tube. Cell debris was pelleted by centrifugation at 21,000 × *g* for 20 mins at 4 °C and the supernatant was aliquoted for use in assays. Crude extract (50 μL) was then added to a new tube containing 27.5 μL reaction buffer (225 mM Tris, pH 7.5, 4.5 mM DTT, 4.5 mM EDTA, 1x proteinase inhibitor cocktail IV), 4.5 μL of 2 mM sulfamethazine (SMZ), and reactions were initiated with the addition of 18 μL of 2.5 mM acetyl-CoA. Negative controls included water in place of acetyl-CoA. Reactions were terminated after 30 minutes by the addition of 50 μL 25% trichloroacetic acid. To each reaction tube, 760 μL of 5% 4-dimethylamino-benzaldehyde in acetonitrile was added and mixed. Tubes were spun for 1 minute at 2,000 × *g* to remove precipitate. Aliquots of 100 μL were then added to a 96-well plate and absorbances at 450 nm were measured. The presence of acetyl-SMZ was determined by loss of signal at 450 nm.

### Growth curves

In brief, individual saturated cultures were prepared for each yeast strain. Cell density was adjusted to 0.8 x 10^7^ cells/mL for all cultures. For single mutants, strains containing pCYP1A2_NAT2 were maintained in selective medium (SC-URA), and exposed in duplicate MeOH, 400 μM IQ, or 800 μM IQ. The plate was covered with optical adhesive tape and incubated at 30 °C in a TECAN Infinite M200 plate reader and absorbance at 600 nm was recorded every 10 minutes for 24 hours, as previously described (Fasullo *et al*., 2010, Fasullo *et al*., 2014, St. John *et al*., 2020). Absorbance was plotted against time and the areas under the curve (AUCs) calculated for the time interval of 0 to 20 hours using freely available software (https://www.padowan.dk/download/). We calculated percent growth in the presence of toxin using the formula (AUC_IQ_/AUC_MeOH_) x 100% (Toussaint *et al*. 2006). Statistical significance of differences between growth percentages for diploid strains and BY4743 were determined by Student’s t-test, assuming equal variance between samples.

### Competitive growth assays

We measured competitive fitness of selected strains using a modified protocol (De la Rosa *et al*., 2007; North *et al*., 2012). In brief, the strain was grown to stationary phase in either YPD or SC-URA (for strains containing pCYP1A2_NAT2). In addition to the strain of interest, a wild-type strain constitutively expressing GFP was grown to saturation (YB675, or YB676 if containing the pCYP1A2_NAT2 plasmid). Cells from the strain of interest and the GFP-expressing strain were combined at a set ratio and inoculated into 2 mL of culture at a concentration of 3×10^5^ cells/mL. These cultures were then treated in duplicate with either solvent, 400 μM IQ, or 800 μM IQ and incubated for 20 hours on a rotary shaker at 30 °C. Cells were then washed thrice with water and resuspended in 1 mL of water. Cells (2×10^4^) were then counted on an Amnis ImageStream^X^ imaging flow cytometer (N = 2). Cells were imaged with brightfield and a 488 nm laser at 50 mW power. The ratio of GFP-expressing cells to total number of cells was compared for solvent-exposed and IQ-exposed cells. Data were analyzed with the IDEAS v6.2 software. In-focus cells were gated, followed by single cells, which included budded cells. GFP fluorescence intensity was quantified for the focused, single-cell population which resulted in two distinct peaks, one for fluorescent cells and one for non-fluorescent cells. Statistical significance of differences was determined by Student’s t-test and Dunnett’s test, assuming constant variance between samples.

### Ames test

To ensure the mutagenicity of IQ, we performed the Ames test with the preincubation protocol (Maron and Ames, 1983). In brief, cultures of *S. typhimurium* TA100 were grown overnight at 37 °C to a concentration of 1-2 x 10^8^ cells/mL. The cultures were placed on ice until preincubation. An S9 master mix was made according to the protocol (33 mM KCl, 8 mM MgCl_2_, 5 mM glucose-6-phosphate, 4 mM NADP, 100 mM phosphate buffer pH 7.4), and to it was added rat liver S9 (4%) or yeast-derived microsomes, and 495 μL of this master mix were added to culture tubes, in triplicate, along with 100 μL of the chilled bacteria. IQ was added at either 4 or 40 μM concentration before incubating the tubes in a heat block at 37 °C for 20 minutes. After incubation, 2 mL of molten 0.6% top agar, heated to 48 °C, was added and the mix was pipetted onto minimal glucose agar plates. Plates were cooled for 5 minutes, then inverted and incubated for 48 hours at 37 °C before His^+^ colonies were counted.

### Trypan blue staining

To measure cell viability after IQ exposure, we performed a trypan blue exclusion assay. Selected strains expressing CYP1A2 and NAT2 were inoculated in SC-URA until cultures reached an A_600_ of 0.1-0.5, and then exposed to either 800 μM IQ or 2% solvent alone. After incubating for 3 hours, cells were washed twice in sterile phosphate buffered saline (PBS) and stained with trypan blue at a final concentration of 2 mg/ml (Liesche *et al*. 2015). Both the total number and blue-stained (dead) cells were counted on a light microscope using a hemocytometer. At least 200 cells were counted for each experiment, and percentages of live and dead cells were calculated, *N* = 3. Statistical significance of percentage differences was determined by Student’s one-tailed *t*-test.

## RESULTS

We profiled the yeast genome for resistance to the heterocyclic aromatic amine IQ using two different pooled collections of the diploid non-essential deletion strains derived from BY4743. The first pool was a yeast deletion collection library of 4607 knock-out strains, each strain containing a single deletion in a non-essential gene (Jo *et al*. 2009). The “humanized” pool expressed CYP1A2 and NAT2 in approximately 4900 strains derived from the 5417 homozygous deletion diploids (St. John *et al*., 2020). Resistance genes were characterized using bioinformatic programs to identify interactomes, protein complexes, and GO process and functional groups. Key genes were then validated.

### Expression of CYP1A2 and NAT2 enhances IQ toxicity in yeast Rad^+^ and *rad4 rad51* mutants

To confirm that expression of CYP1A2 and NAT2 bioactivate IQ, the expression plasmid pCYP1A2_NAT2 was introduced in wild type yeast (BY4743) and a *rad4 rad51* strain by selecting for Ura^+^ transformants. Expression of CYP1A2 and NAT2 proteins was then confirmed by Western blots (Figure 2E and 2F, respectively). We performed MROD assays to quantify CYP1A2 activity (Figure S1A, Fasullo *et al*., 2014) and N-acetyl transferase (NAT) assays to detect NAT2 activity (Figure S1B). These extracts also activated IQ in the Ames assay; compared to the strain expressing no human genes, the mutagenicity of IQ was highest (P < 0.05, Table S3) when IQ was preincubated using microsomal extracts obtained from strains expressing CYP1A2 and NAT2, but still detectable using extract from strains expressing CYP1A2 alone (Table S3). These data indicate that CYP1A2 and NAT2 were expressed in yeast and were sufficient to increase the mutagenicity of IQ.

**Figure 2.**
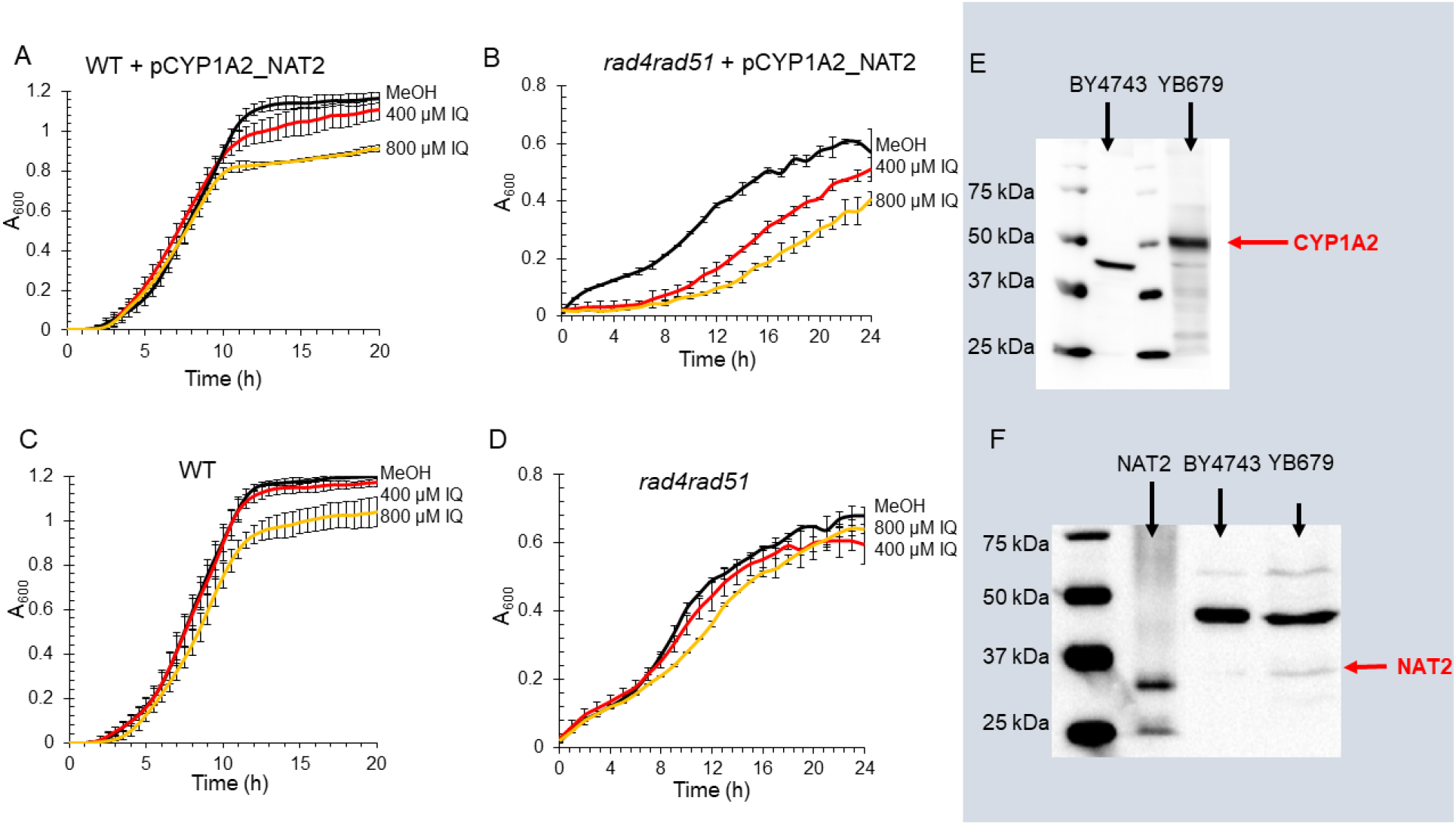
IQ sensitivity and expression of CYP1A2 and NAT2 in Rad^+^ and DNA repair deficient strains. Panels A, B, C, and D show growth curves of Rad^+^ or *rad4 rad51* yeast strains exposed to 2% MeOH (black), 400 μM IQ (Red), or 800 μM IQ (Orange), plotted as absorbance (A_600_) against time (Hs). Error bars represent one standard deviation at half-hour intervals. Panel **A** is a growth curve of the wild-type diploid strain containing pCYP1A2_NAT2 (YB679), N = 8. Panel **B** is a growth curve of the *rad4 rad51* haploid yeast strain containing pCYP1A2_NAT2 (YB400), N = 3. Panel C is a growth curve of the Rad^+^ diploid strain (BY4743), N = 8. Panel D is a growth curve the *rad4 rad51* (YB226) strains, N =3. Panel **E** is a Western blot that detects the human CYP1A2 protein present in yeast protein extracts. The red arrow indicates the position of CYP1A2, which is ~58 kDa. The first and third lanes from the left contain the molecular weight standards. The second lane contains the protein extract from wild type (BY4743) and the last lane contains the extract from wild-type strain containing pCYP1A2_NAT2 (YB679). Panel **F** is a Western blot that detects the human NAT2 protein in yeast protein extracts. The red arrow indicates the position of the NAT2 protein, which is ~37 kDa. The first lane from the left contains the molecular weight standards. The second contains a purified NAT2 protein. The third and fourth lanes respectively contain the wild-type (BY4743) and the wild type expressing pCYP1A2_NAT2 (YB679).

By calculating areas under the curve (AUCs), we quantified IQ concentrations that reduced growth in both the wild-type Rad^+^ diploid (BY4743) and the *rad4 rad51* haploid mutant, which either did (“humanized”) or did not express CYP1A2 and NAT2. Exposure to 400 μM IQ and 800 μM IQ reduced growth in the Rad^+^ diploid by 1% and 7%, respectively, and in the *rad4 rad51* haploid by 6% and 13%, respectively. On the other hand, exposure to 400 μM IQ and 800 μM IQ, reduced the growth of the “humanized” Rad^+^ diploid by 6% and 17%, respectively, and in the “humanized”*rad4 rad51* haploid by 40% and 59% (Figure 2). The data indicate that expression of CYP1A2 and NAT2 bioactivates IQ into a yeast genotoxin.

### Identification of resistance genes by barcode analysis

We identified genes that confer IQ resistance at two different time exposures and after two different IQ concentrations, for a total of six screens (Figure 3). We exposed the pooled “humanized” yeast diploid collection after 800 μM IQ exposure for 5G and 10G, and then independently at 400 μM and 800 μM exposures for 10G. We exposed the original yeast deletion library at 400 μM and 800 μM exposures for 10G. The control for all screens was exposed to 2% methanol and four biological replicates were performed for each exposure. Approximately 98% (4513/4607) of strains from the original library and 89% (4343/4900) of strains from the humanized library were detected in the pooled cultures.

**Figure 3:**
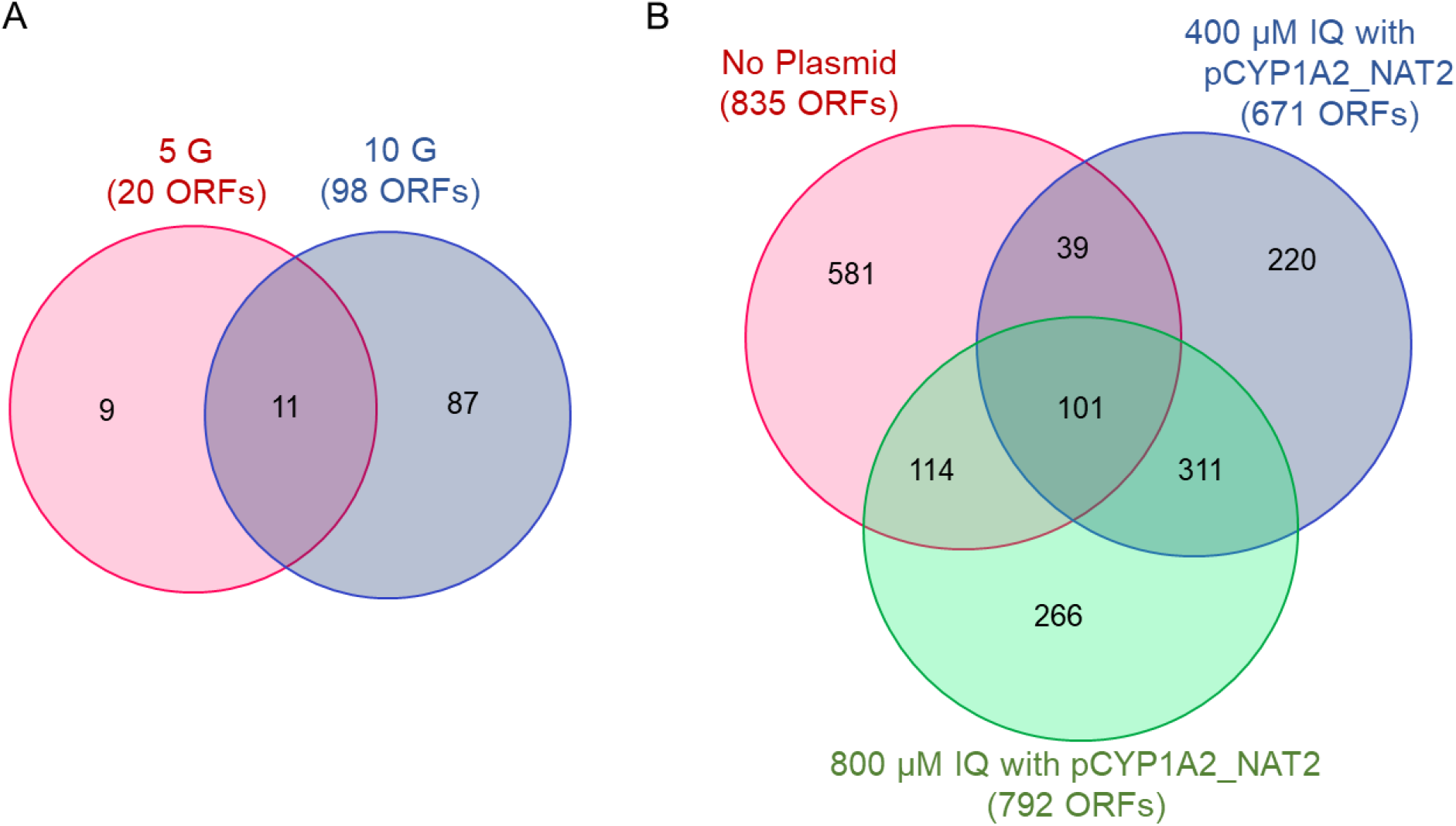
Venn diagrams of IQ resistance genes identified in screens that vary either exposure time or IQ concentrations. All ORFs were identified from the yeast diploid deletion library expressing CYP1A2 and NAT2. Panel A shows the overlap of the IQ resistance ORFs identified after 800 μM IQ exposure for five and ten generations. 20 resistance ORFs were identified after five generations and 98 ORFs were identified after ten generations. Panel B shows the overlap of the IQ resistance genes identified after 400- and 800- μM IQ exposures for ten generations. Individually, 671 ORFs were identified after 400 μM IQ exposure and 792 ORFs were identified after 800 μM IQ exposures. The original library treated with either IQ concentration identified 835 ORFs. ORFs include genes of known function and uncharacterized open reading frames.

We classified genes that confer IQ resistance (m < 0, q < 0.1) as 1) those that confer resistance to IQ without CYP1A2 and NAT2 activation, and 2) those that confer IQ resistance after CYP1A2 and NAT2-mediated activation. In total, 1159 ORFs from the “humanized” library were identified that conferred IQ resistance (see Table S4), of which, 337 genes of known function were identified in at least two independent screens or were validated (see Table S5). Strains identified that conferred IQ resistance in all four screens of the “humanized” collection were *CSG2, RAV2, HCR1*, and *SFG1*. From the original library, 835 resistance ORFs were identified from exposures at either 400 μM and 800 μM IQ (Table S6). Of these, 143 genes of known function conferred resistance at both IQ concentrations (Table S7). 101 ORFs were identified that conferred IQ resistance from both exposures of the “humanized” and at least one exposure of the “original” libraries (Table S8). The Venn diagram with the numbers of significant IQ resistance ORFs from these experiments and their overlap is presented in Figure 3.

### Interactome of IQ resistance genes from library expressing CYP1A2 and NAT2

To validate key pathways that mediate IQ resistance in strains expressing CYP1A2 and NAT2, we identified a network of gene interactions using ELIXIR’s Core Data Resource STRING (https://string-db.org/, Szklarczyk *et al*. 2018) and protein complexes, using Princeton GO Term finder (https://go.princeton.edu/cgi-bin/GOTermFinder, Boyle *et al*., 2004). An interactome for proteins encoding IQ resistance genes from the original library is shown in Figure S2. The network of protein interactions of all significant IQ resistance ORFs from both 400 μM IQ and 800 μM IQ treatments is shown in Figure 4 (left panel). The cluster that has the most members are ribosomal proteins. Other clusters include kinases, phosphatases, chaperones and thermotolerance proteins and proteins that participate in DNA damage repair, nitrogen metabolism, and histone modification.

**Figure 4:**
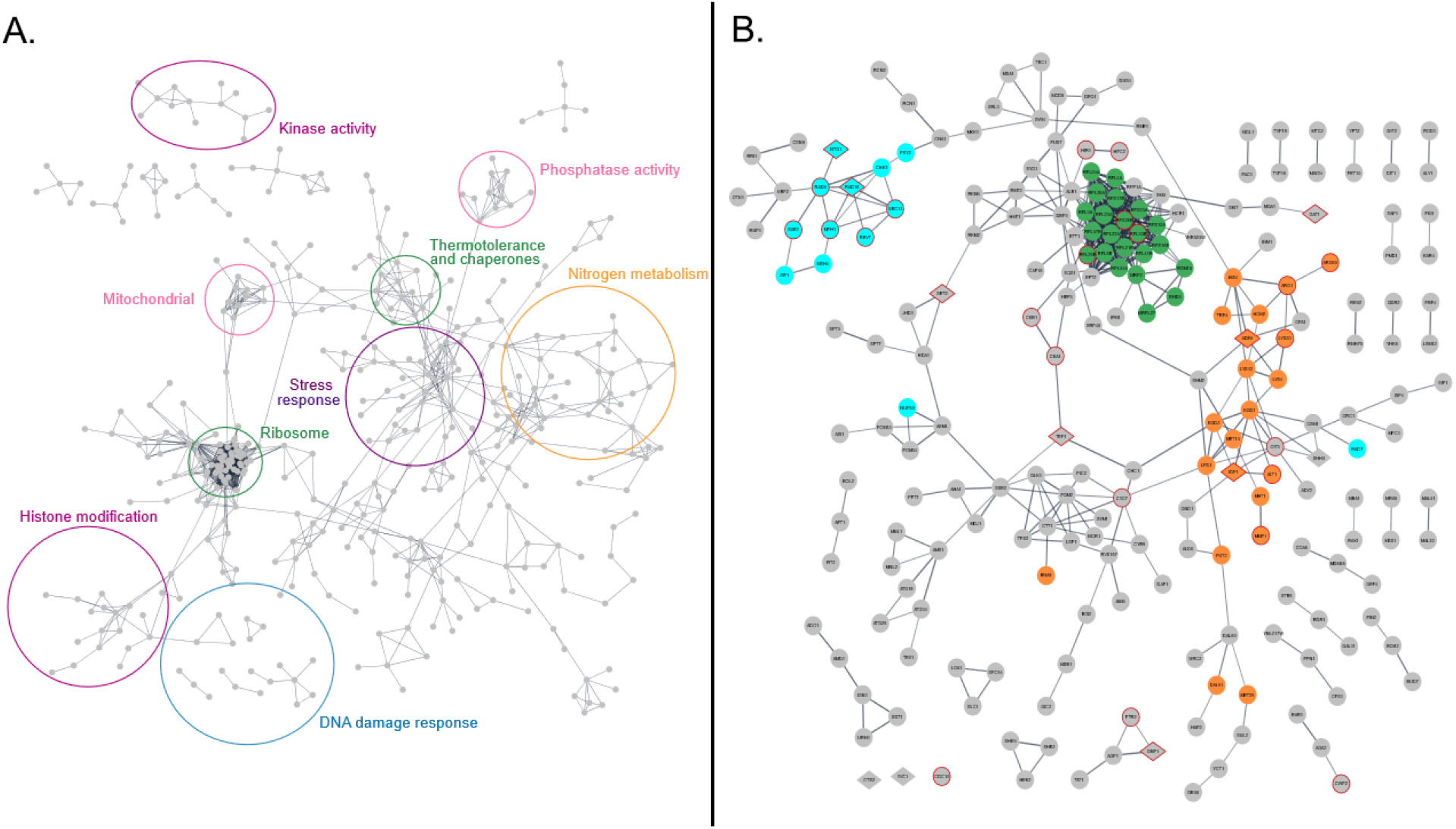
Protein interactome identified from screens of the diploid deletion library containing pCYP1A2_NAT2. The interactome was curated using String V11 (https://string-db.org, Szklarczyk *et al*. 2019). Panel A represents the functional interactome of each protein encoded by ORFs that were identified in at least one screen, using the highest confidence level of 0.9. Panel B represents the functional interactome of each protein encoded by genes identified from at least two screens of the yeast deletion library containing pCYP1A2_NAT2, using a high confidence level of 0.7. The *rad9, rad18, mph1, and ntg1* strains were independently validated (Figure S3 and S4) and included in Panel B. Colored nodes indicate protein translation (green), DNA damage tolerance (blue), and amino acid metabolism (yellow). Genes with human orthologs implicated in colon cancer are shown as diamond-shaped nodes. Nodes circled in red were validated. The unconnected nodes shown in the bottom left were either individually validated or are implicated in colon cancer. The rest of the unconnected notes were removed to improve readability.

To focus on specific genes that participate in these pathways, we constructed an interactome for proteins encoded by IQ resistance genes that were significant in at least two different screens of the “humanized” library (Figure 4B). The largest and most heavily connected cluster was ribosomal proteins. Additional clusters are encoded by DNA damage tolerance genes, such as *REV7, UBC13*, and *RAD18*, and nitrogen metabolism genes, such *ALT1, ADE6, LYS20*. Proteins encoded by several genes are responsive to nitrogen stress, such as *RME1* and *TEC1*, and general nutrient stress, such *GUF1* and *RPH1*. These data thus confirm several key gene interaction networks involved in protein metabolism and DNA damage repair.

Protein complexes in global cell regulation and housekeeping were identified among the 337 genes (P <0.05), and redundancy was reduced using Revigo (Supek *et al*., 2011). The resulting 14 complexes included the integral component of nuclear outer membrane, ribosome, the protein kinase CK2 complex, the HIR complex, Cvt complex for autophagy, and the spindle complexes (Table 1); the nuclear outer membrane complex was the most highly enriched (13-fold) complex. Both the CK2 kinase, nuclear membrane, and spindle complexes regulate cell cycle progression, while the HIR complex, also referred to as the replication-independent histone chaperone complex, regulates chromatin structure and affects gene expression. The Cvt complex is important in protein degradation and indicates that removal of damaged proteins is important in IQ resistance. These complexes underscore the importance of regulating cell cycle progression and protein turnover in IQ resistance.

**Table 1:** Protein complexes that confer IQ resistance

| Term ID | GO Group | Number in set<sup>1</sup><br>(Cluster %) | Number in Library<sup>2</sup><br>(Background %) | Enrichment<sup>3</sup> | P-Value | Genes |
| --- | --- | --- | --- | --- | --- | --- |
| GO:0031309 | Integral component of nuclear outer membrane | 2 (0.59%) | 2 (0.046%) | 13 | 6.2E-03 | *POM34, POM33* |
| GO:0034270 | Cvt complex | 3 (0.88%) | 4 (0.093%) | 9.5 | 1.8E-03 | *ATG19, ATG34, AMS1* |
| GO:0001400 | Mating projection base | 2 (0.59%) | 3 (0.069%) | 8.5 | 1.8E-02 | *AFR1, CDC10* |
| GO:0045240 | Dihydrolipoyl dehydrogenase complex | 3 (0.88%) | 5 (0.12%) | 7.6 | 4.3E-03 | *KGD1, KGD2, LPD1* |
| GO:0000417 | HIR complex | 2 (0.59%) | 4 (0.093%) | 6.4 | 3.3E-02 | *HPC2, HIR1* |
| GO:0005956 | Protein kinase CK2 complex | 2 (0.59%) | 4 (0.093%) | 6.4 | 3.3E-02 | *CKA1, CKB1* |
| GO:0034456 | UTP-C complex | 2 (0.59%) | 4 (0.093%) | 6.4 | 3.3E-02 | *CKA1, CKB1* |
| GO:0045239 | Tricarboxylic acid cycle enzyme complex | 3 (0.88%) | 8 (0.19%) | 4.8 | 2.0E-02 | *KGD1, KGD2, LPD1* |
| GO:0042645 | mitochondrial nucleoid | 4 (1.2%) | 17 (0.39%) | 3.0 | 4.0E-02 | *KGD1, KGD2, IDP1, LPD1* |
| GO:0005819 | spindle | 6 (1.8%) | 29 (0.67%) | 2.6 | 2.3E-02 | *SLI15, PAC1, BFA1, SPO13, MPC54, CDC10* |
| GO:0005840 | ribosome | 25 (7.4%) | 186 (4.3%) | 1.7 | 5.2E-03 | *RPS16B, RPL22A, RSM18, RPL33B, RPL23A, RPP1A, RPL39, EHD3, RPS17B, BTT1, RPS10A, RPL26B, HEF3, RPL24A, MRP2, RPL6B, RPL37B, RPL4A, RPL34A, RPL21B, RPL41B, MRPL27, RPL26A, RPS26B, RPS23A* |
| GO:1990904 | ribonucleoprotein complex | 32 (9.4%) | 270 (6.3%) | 1.5 | 1.1E-02 | *SQS1, RPL22A, RPL33B, RPL23A, RPL39, EHD3, BTT1, RPL24A, MRP2, RPL37B, ESL2, RPL4A, RPL34A, MRPL27, MNE1, RPS26B, RPS23A, RPS16B, RSM18, RPP1A, RPS17B, RPL26B, RPS10A, HCR1, DBP3, CKA1, RPL6B, CKB1, EFT2, RPL21B, RPL41B, RPL26A* |

<sup>1</sup>: 337 genes in set<sup>2</sup>: 4343 genes in background<sup>3</sup>: Enrichment = Cluster% / Background%

We identified 18 functional GO groups represented by the 337 genes (P < 0.05) using Princeton GO Term finder and reduced redundancy using Revigo (Table 2). While the ribosome cluster was again prominent, other GO terms included phosphatase activity, quaternary ammonium group transmembrane transporter activity, sulfur compound transmembrane transporter activity, DNA and RNA binding, and purine ribonucleotide transmembrane transporter activity. These process terms emphasize the role of transport in conferring IQ resistance, as well as the role of nucleic acid binding.

**Table 2.**
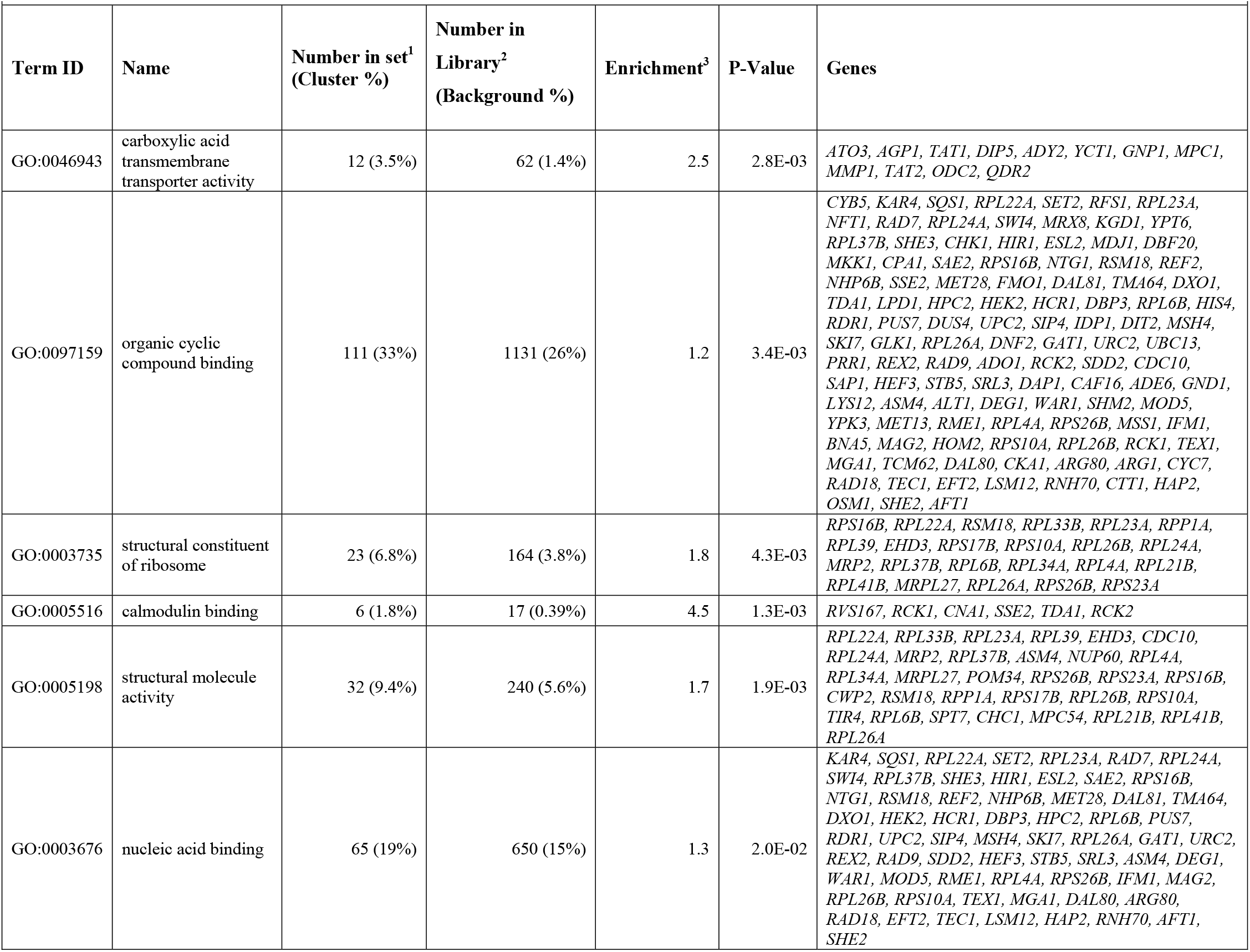
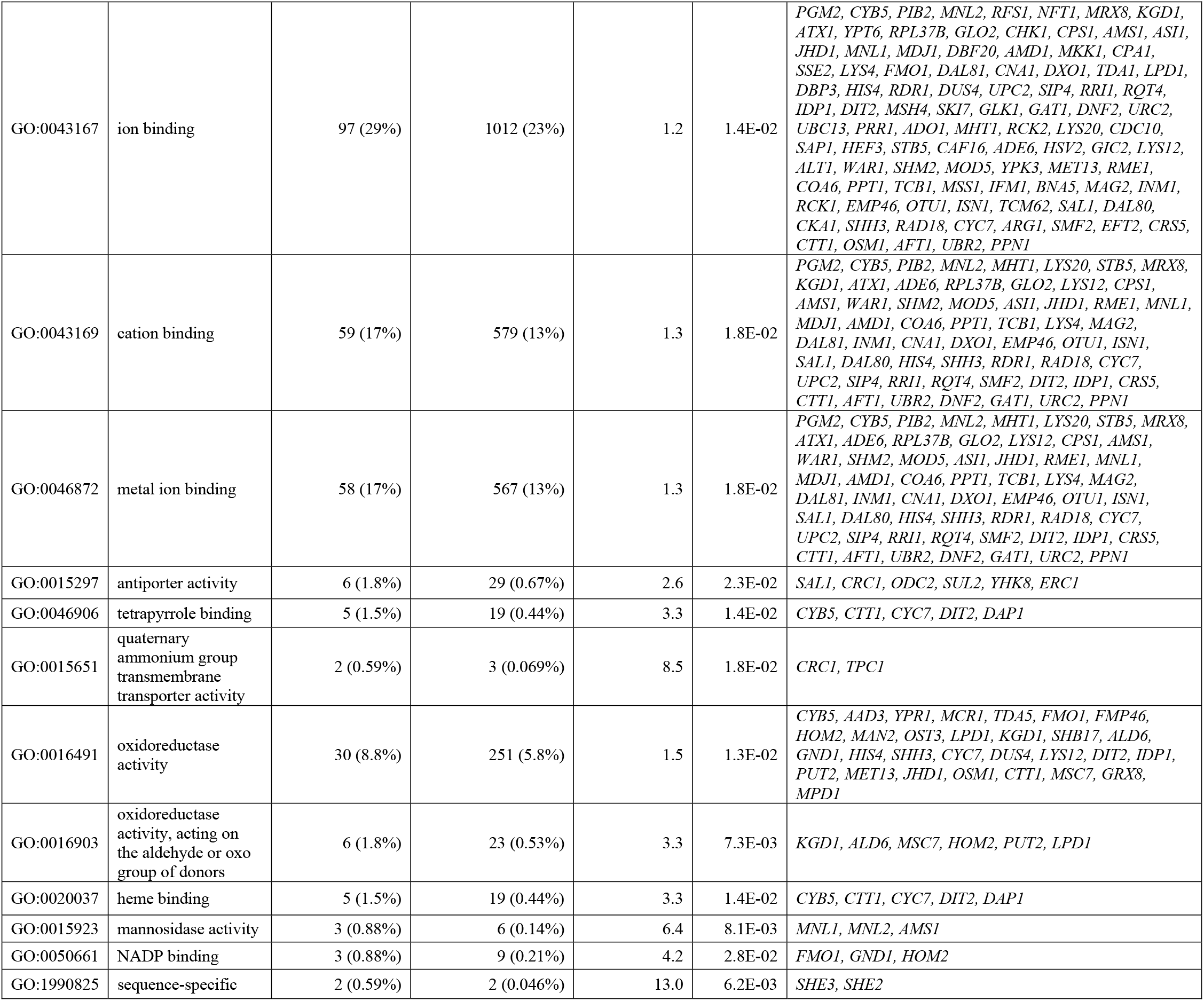
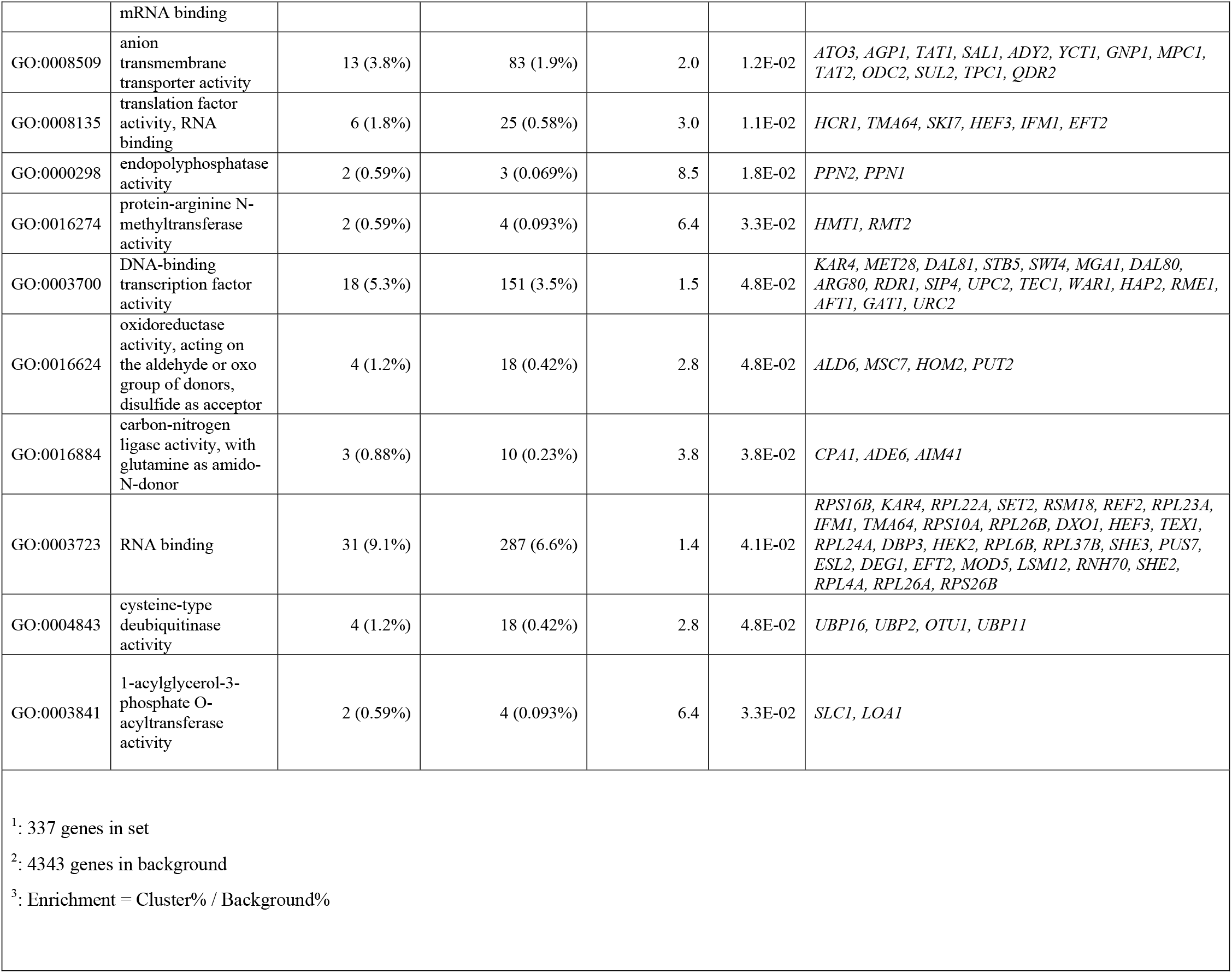
GO Function Terms for validated IQ resistance genes from humanized library

We listed 22 IQ resistance genes involved in either maintaining genome stability, DNA repair, or DNA damage-associated gene expression (Table 3). While several genes were directly associated with DNA damage tolerance, such as *UBC13, REV7*, and *RAD18*, others participated in cell cycle progression in the presence of DNA damage, including *PSY2, CKA1* and *CKB1. ADE6*, which functions in *de novo* purine synthesis, suppresses chromosome loss, as revealed in a *MAT**a***-like faker screen (ALF, Yuen *et al*., 2007). Recombinational repair genes included *MPH1, SAE2*, and *IES4*. Checkpoint signaling genes included *RAD9* and *CHK1. NUR1* functions in maintaining rDNA structure. While NER genes were not prominent, we identified the dual function base excision repair gene, *NTG1*. These genes emphasize the role of DNA damage tolerance in conferring IQ resistance.

**Table 3:**
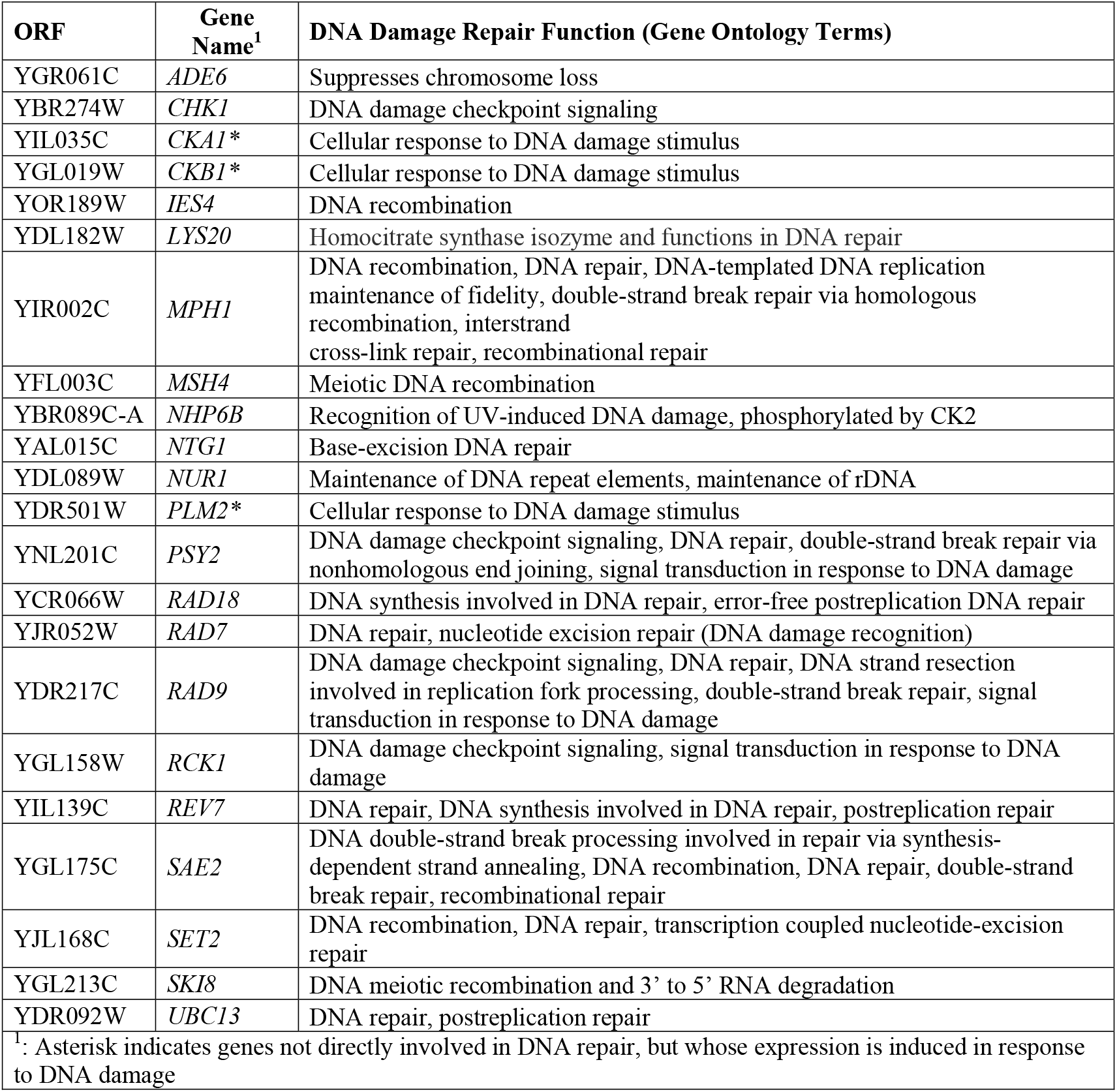
Genome stability, DNA damage repair and DNA damage-inducible genes validated or identified in multiple IQ resistance screens

Human homologs of IQ resistance genes included 11 genes associated with colorectal cancer and ten genes associated with other or multiple cancers (Table 4). Biallelic mutations in the human ortholog of *NTG1, NTHL1*, are a high-risk factor for familial adenomatous polyposis (Groden *et al*., 1991). The human homolog for *RAD18* has been noted to participate in multiple cancer progression stages and over-expression confers resistance to chemotherapeutic agents (Li *et al*., 2020, Yang *et al*., 2018). Alleles in other colorectal cancer risk-associated genes include *SET2* and *SHH3* (Choi *et al*., 2014, Habano *et al*., 2003). In addition, other genes were associated with multiple cancers, including breast and colon cancer. These include *ADE6*, which encodes phosphoribosylformylglycinamidine (FGAM) synthase, and *TEP1*, which is the yeast ortholog for the tumor suppressor gene *PTEN*.

**Table 4:**
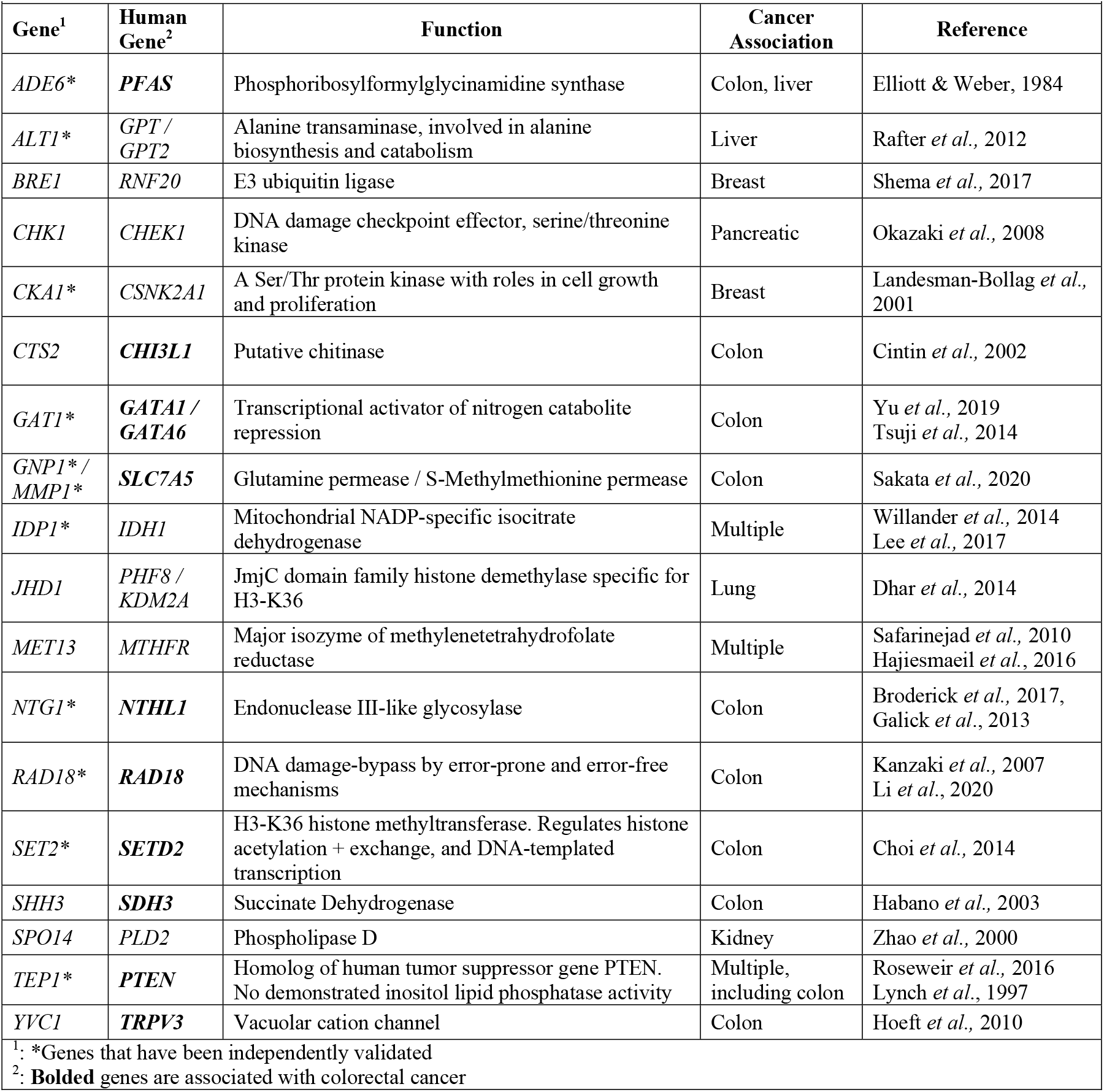
Yeast genes conferring resistance to activated IQ with human orthologs associated with cancer

### Validation of Genes

We focused on validating IQ resistance yeast genes relating to colon cancer and identified in the major interactome clusters using growth curves, competitive growth assays, and trypan blue staining; while growth curves and competitive growth assays measure toxin resistance after 10g exposures, trypan blue was used to determine cell viability after short-term 3h IQ exposure. We previously observed that many strains lose the CYP1A2 expression plasmids rendering it difficult to accurately determine toxin sensitivity by growth curves (St. John *et al*., 2020). In total, we validated 38 out of 51 selected genes (75%), of which 12 were validated by multiple methods, and three (*ALT1, ADE6, RAD18*) were validated by all three methods. Representative strains from each of the highlighted clusters of the interactome are shown in Figure 5, along with the “wild-type” strains.

**Figure 5.**
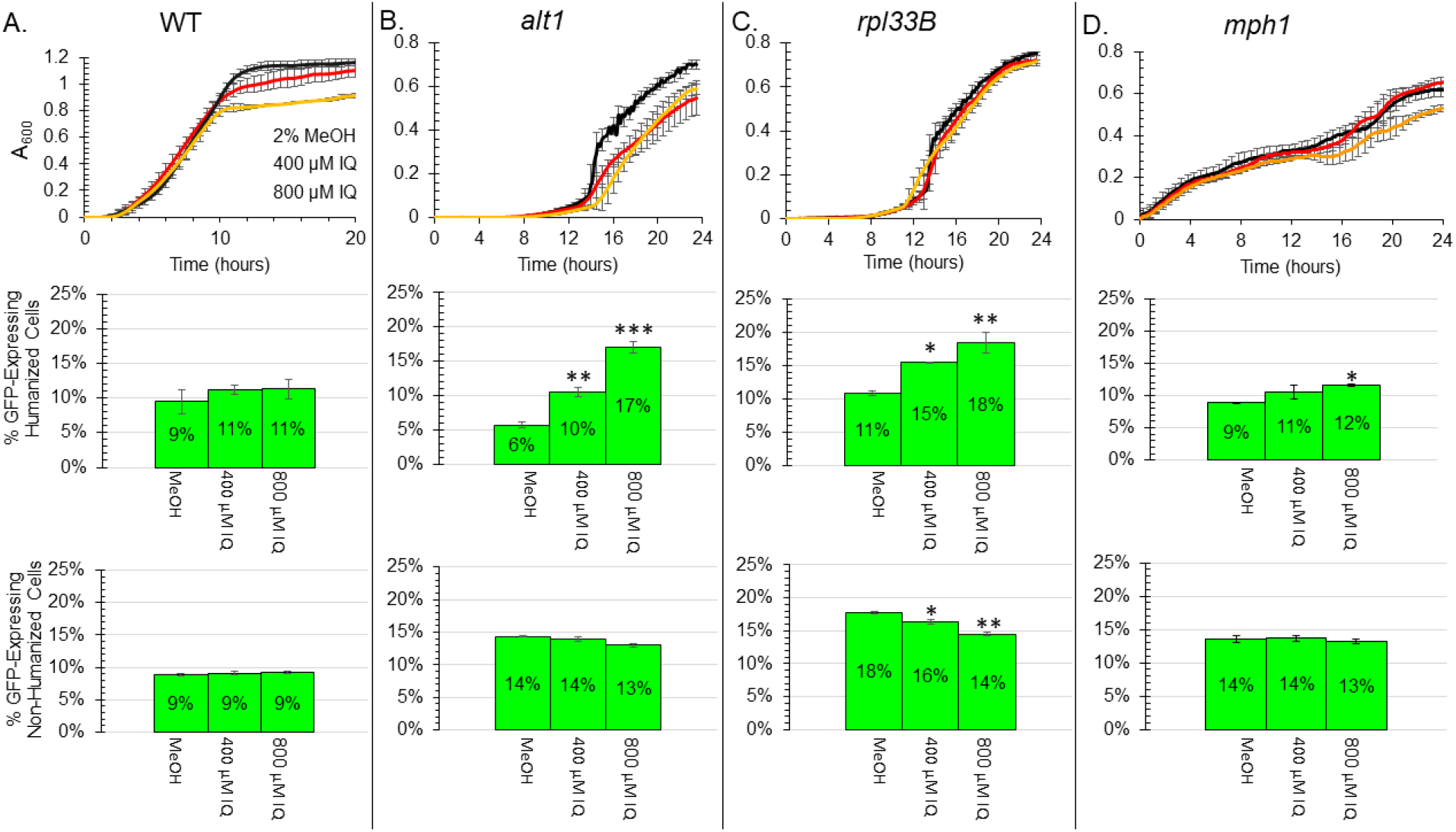
Growth curves and competition assays for selected IQ resistance genes identified in high-throughput screens. Panels A, B, C, and D show the wild-type strain, *alt1, rpl33B*,and *mph1*, respectively. The latter three represent major gene clusters highlighted in Figure 4, including nitrogen metabolism (*ALT1*), ribosome protein (*RPL33B*) and DNA repair (*MPH1*).Columns **A, B, C,** and **D** are wild type, *alt1, rpl33B*, and *mph1* diploid strains, respectively. The top row contains growth curves for selected strains containing pCYP1A2_NAT2. A_600_ is plotted against time (Hs) with the background absorbance was subtracted, N > 3. Error bars represent one standard deviation for half-hour measurements. The middle row shows competitive growth data for the indicated strains containing pCYP1A2_NAT2. The y-axis shows the percent of GFP-containing wild type expressing CYP1A2 and NAT2 (YB676). Competitive cultures were inoculated into SC-Ura at approximately 10% YB676 and 90% the indicated strain. 2×10^4^ cells were acquired per sample. Error bars represent one standard deviation. For wild type (YB679), N = 5; for all others, N=2. Significance was determined by Dunnett’s test and asterisks indicate p<0.05 (*), p<0.01 (**) and p<0.001(***). The bottom row shows competitive growth data for the indicated strains not expressing CYP1A2 or NAT2. The y-axis shows the % of GFP-containing WT (YB675). Competitive cultures were inoculated into SC at approximately 10% YB675 and 90% the indicated strain. 2×10^4^ cells were acquired per sample. For wild type (BY4743), N=4; for all others, N=2. Error bars represent one standard deviation. Significance was determined by Dunnett’s test and asterisks indicate p<0.05 (*), p<0.01 (**) and p<0.001(***).

Mutants *alt1, rpl33B, mph1* were selected because they map to nitrogen, ribosome, and DNA repair clusters (Figure 4). They exhibited particularly high IQ sensitivity either by growth curves or competitive growth assays. *ALT1* functions in utilizing alanine as a nitrogen source, and *alt1* mutants are sensitive to oxidative stress. *RPL33B* encodes the ribosomal 60S genes and which is essential when its paralog *RPL33A* is defective. Other genes that were validated in these clusters *RPL23A, RPL23B, LYS20, ARG1, RAD18, RAD9, REV7*, and *SAE2* (Tables S4 and S5, Figure S3, S4, and S5).

We quantified IQ-associated lethality in mutants defective in candidate resistance genes. Percentage of dead cells after treatment with solvent or IQ are shown in Figure S4 for all strains tested. BY4743 was used as the wild-type strain for diploids, while BY4741 was used for the wild-type control in haploids. Mutants that exhibited a high percentage of IQ-associated death included *cdc10, arg1, arg80, cka1*, and *ubc13*, while the *rad18* mutant exhibited a five-fold increase in percent of dead cells after IQ exposure.

### Enhanced sensitivity of haploid double mutants defective in DNA repair

We hypothesize that one reason we did not identify more DNA repair genes is that there are multiple repair functions for IQ-associated DNA damage. To assess this hypothesis, we made *rad4 rad51, rad4 rad18, rad4 ntg1*, and *rad18 rad51* haploid double mutants. We compared the corresponding single and double mutants using growth curves, competitive growth, and viability assays. Competitive growth data for these double mutants, along with the haploid single mutants, expressing CYP1A2 and NAT2 are seen in Figure 6. IQ-associated cell death data are shown in Figure S4. Growth curves for double and single mutants are shown in Figure S5. Compared to respective single mutants, the IQ sensitivity was most enhanced for the *rad4 ntg1* and *rad4 rad51* double mutants. Collectively these data suggest that *NTG1* and *RAD4*, and *RAD4* and *RAD51* function in multiple DNA repair pathways for IQ-associated DNA damage.

**Figure 6:**
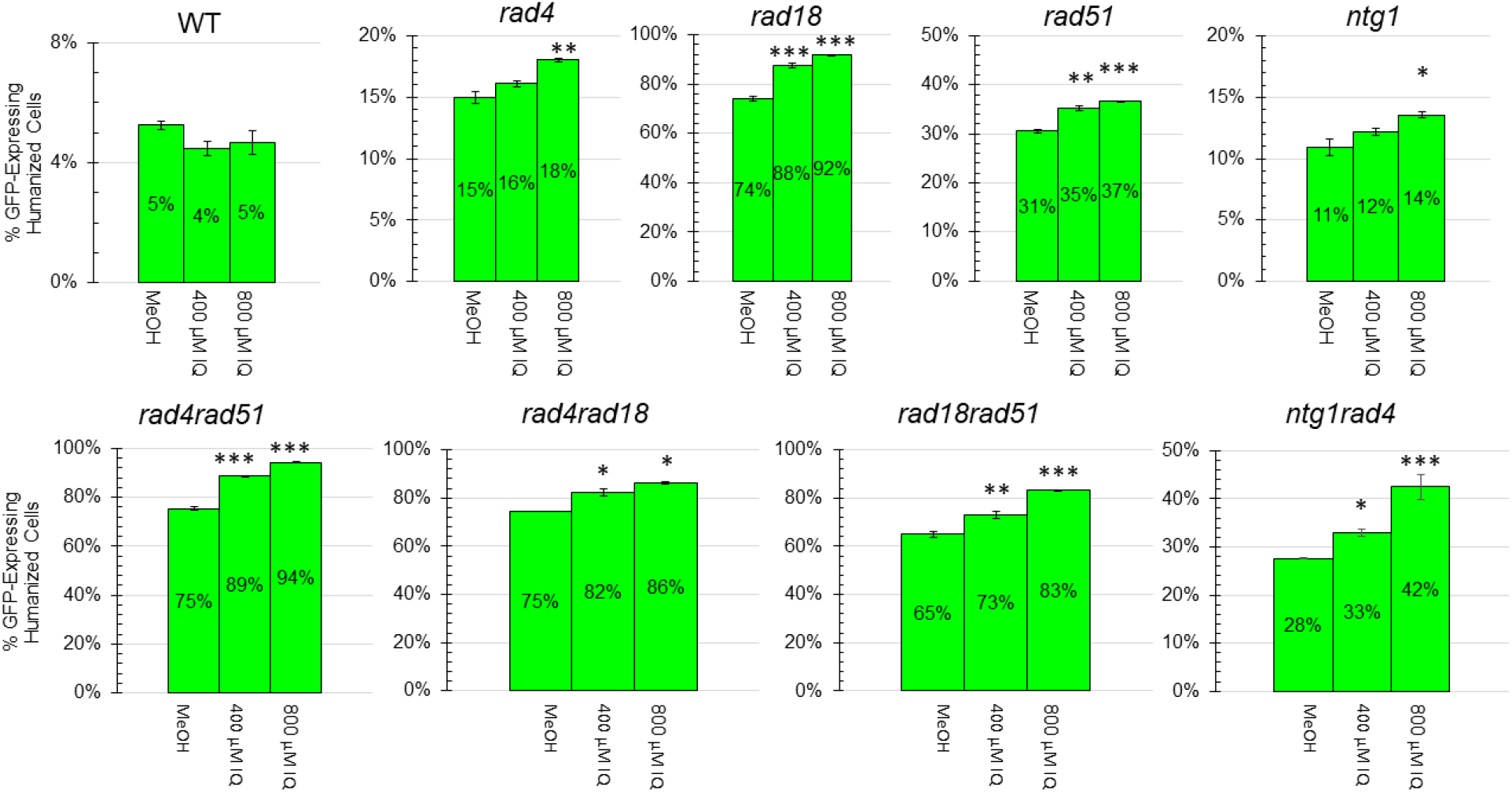
IQ toxicity in *rad4, ntg1, rad18*, and *rad51* single in double mutants. The competitive growth of selected single haploid DNA repair mutants (top row) and double mutants (bottom row) after growth in the presence of 2% MeOH, 400 μM IQ, or 800 μM IQ. Cultures were inoculated at a ratio of 10% of GFP-expressing wild-type cells to 90% mutant cells and were grown for 20 hours (~10 generations). 2 x 10^4^ cells were acquired for each sample. For the WT haploid (YB682), N=8. For the other strains, N=2. Error bars represent one standard deviation. Significance was determined by Dunnett’s test and asterisks indicate p<0.05 (*), p<0.01 (**) and p<0.001(***).

### IQ resistance genes from original yeast library

The protein interactome of all 835 IQ resistance ORFs identified from the original collection is shown in Figure S2A and the interactome derived from 143 ORFs identified in at least two screens is shown in Figure S2B. In Figure S2A the ribosomal cluster is clearly visible; four genes, *RPL22A*, *RPL26A, RPL37B*, and *RPL6B*, were identified in profiling both the “humanized” yeast library and the original library, however, only *RPL22A* appeared in at least two screens of each library. Among GO terms over-represented were negative regulation of nucleobase-containing compound metabolic process (GO:0045934) and cellular protein modification process (GO:0006464). There were 101 genes that appeared in both “humanized” library and at least one screen of the original yeast library (Figure 3). While 19 out of the 101 genes that conferred IQ resistance from both the “humanized” and the original library have no known function, genes related to chemical accumulation include *RDR1*, which is a negative regulator of *PDR5* and whose over-expression increases drug resistance (Hellauer *et al*., 2002), and *OSM1*, which confers osmotic resistance (Singh and Sherman, 1978). Other genes included *CTT1*, which encodes catalase and confers resistance to oxidative stress (Lushchak and Gospodaryov, 2005), and *DAL80*, which is a negative regulator of nitrogen catabolism genes (Marzluf, 1997). These studies suggest that in budding yeast, IQ exposure *per se* disrupts nitrogen metabolism and causes oxidative stress.

We also identified IQ sensitive genes (m > 1, q < 0.1) that are not included in the Venn Diagram. These are listed in Table S9. Among these, there were a few that were initially identified as IQ resistance genes, including *NTG1* and *RAD18;* however, in validation assays these genes conferred resistance. Sensitive genes include those that function in nitrogen compound and organic substance transport; we speculate that transporting IQ into cellular compartments may enhance sensitivity. Genes that confer IQ sensitivity may also stabilize the pCYP1A2_NAT2 plasmid, thus increasing IQ activation.

## DISCUSSION

IQ is a class 2A and rodent carcinogen, present in charred meats in cigarette smoke. In this study we profiled the yeast genome for IQ resistance by screening a “humanized” yeast diploid deletion library that expresses both CYP1A2 and NAT2. We identified 337 resistance genes that appeared in at least two of four independent screens, and validated 38 using either growth curves, competitive growth assays, or viability assays. Several are responsive to DNA replication stress (30/337) or oxidative stress (9/337), and either their expression is increased, or the encoded proteins are relocated. Major ontology groups included DNA binding, phosphatase, and ribosomal protein genes, while protein complexes included HIR2 and CK2. Human orthologs of 11 yeast genes are associated with colorectal cancer. We speculate that the colon cancer risk associated with these genetic factors could be increased by exposure to HAAs.

Three conclusions can be drawn from our studies screening the “humanized” and original yeast library. First, both screens identified ribosomal protein genes as resistance genes, thus suggesting that IQ targets ribosomes, and by extension, protein synthesis. Second, prominent among DNA repair genes were post-replication repair genes and *NTG1*, while NER genes were less prominent. Third, a comparison of the toxicity of IQ with other toxins, such as AFB_1_ (Table S11), underscores the role of important genes and complexes involved in DNA damage and genotoxin tolerance, including *RAD18* and those encoding the CK2 complex.

These conclusions were based on profiling the yeast genome for IQ resistance using a “humanized” library that expresses both CYP1A2 and NAT2. Nonetheless, there were limitations to this study. First, some DNA repair deficient strains were not detected even after exposure for five generations; a similar limitation was observed in a screen of the “humanized” library for AFB1 resistance and was partially attributed to CYP1A2 expression, which confers an enhanced growth disadvantage (St John *et al*., 2020). For example, *mre11, xrs2*, and *rad52* strains were not detected in our screens. Second, genome profiling necessitated IQ concentrations that likely conferred toxicity by additional nongenotoxic mechanisms. While IQ sensitivity is enhanced in strains expressing CYP1A2 and NAT2, we do not know the exact proportion or amounts of activated IQ metabolites and all their associated targets.

Among the ribosomal protein genes required for resistance to both “activated” IQ and IQ, the oxidative stress response gene, *RPL22A* (Chan *et al*., 2012), is present in both sets, while genes that function in rRNA export from the nucleus and genes encoding proteins with KOW motifs were only present among “activated” IQ resistance genes. Both IQ and activated IQ resistance genes encode rRNA proteins located in the large and small ribosomal subunits; however, there are more “activated” IQ resistance genes that function in a wider set of RNA translation functions and ribosomal protein transport (Table S4, Table S6, Figure 4A and S2A). This set included genes that facilitate RNA translation, such as *DEG1*, modify tRNAs, such as *PUS7, MOD5* and *DUS1*, and import ribosomal proteins into the nucleus, such as *SYO1* (Table S4 and Figure 4A). Taken together, these data suggest that in comparison to resistance to IQ, resistance to activated IQ requires a larger set of genes involved in protein translation and rRNA transport.

Many ribosomal protein genes that confer IQ resistance also confer resistance to other toxins. Typical cellular reactions to multiple stresses involve downregulation of ribosomal protein gene transcription (Warner, 1999; Gasch *et al*., 2000), which may render ribosomal protein mutants sensitive to mycotoxins and antifungal agents. Ribosome quality control is a central feature of resistance to multiple toxins, including trichothecene mycotoxins and cycloheximide (Kugler *et al*., 2016). Nonetheless, major cycloheximide resistance genes, *RPL41, RPL42* and *RPL29* (Dehoux *et al*. 1992; Bae *et al*., 2018; Käufer *et al*., 1983) did not appear in our screen for IQ resistance but instead appeared to confer IQ sensitivity (Table S9).

The notion that activated IQ targets proteins is supported by observations that human serum albumin (SA)Cys^34^ readily reacts with HAAs (Turesky and Le Marchand, 2011; Bellamri *et al*., 2021) and that IQ reacts with heme containing proteins (Turesky *et al*., 1987), suggesting that similar reactions could occur with yeast proteins. IQ-associated protein modification may explain why ammonia, amino acid metabolism, and protein synthesis were heavily represented among HAA resistance genes (Table 2 and S10). Since protein modification genes (*ALG3, ALG5, ALG6, ALG8, ALG12*) were also identified in the IQ screen of the original deletion library (Table S6, Figure S2), we presume that some cellular damage could also occur by IQ exposure *per se* or by IQ metabolites generated by endogenous yeast enzymes.

Our data supports the notion that replication bypass functions in IQ resistance, as was suggested by *in vitro* experiments using oligonucleotides containing IQ-N^2^ Gua adducts (Stavros, 2014). DNA repair genes that conferred IQ resistance were mostly confined to those participating in post-replication repair. These included *RAD18*, *REV7*, and *RAD9*, which participate in DNA damage tolerance pathways that bypass DNA damage by error-prone polymerases (for review, see Xu *et al*., 2015). We also identified *MPH1*, a gene whose helicase function resolves stalled DNA structures, and which functions in error-free repair (Schürer *et al*.,2004).

*RAD18* and *REV7*, which confer IQ resistance, are also common resistance genes for multiple cross-linking agents and agents that cause DNA adducts. These include trichloroethylene (De la Rosa *et al*., 2017) and AFB_1_ (Table S11, St. John *et al*., 2020), as well as chemotherapeutic drug derivatives, such as cisplatin, oxaliplatin and mitomycin C (Wu *et al*.,2004). *RAD18* may also function in error-free mechanisms that efficiently bypass N^2^-Gua adducts (Washington *et al*., 2002). The contribution of both error-free and error-prone mechanisms in bypassing IQ-associated DNA adducts will require additional studies.

Other protein kinases and phosphatases may have secondary roles in conferring tolerance to IQ associated damage. Both the CK2 complex genes and *PSY2* confer tolerance to multiple DNA damaging agents (St. John *et al*., 2020). *PSY2* encodes phosphatase that deactivates Rad53 after exposure to a subset of DNA damaging agents (O’Neill *et al*., 2007), while the CK2 complex is required for toleration of irreparable double-strand breaks (Toczyski *et al*., 1997).

In the repair of IQ-N^2^ Gua or oxidized DNA bases, NER may not serve as a primary DNA repair mechanism, as suggested by Stavros *et al*., (2014). This may partially explain why *rad4* mutants exhibited modest IQ sensitivity, while the double *rad4 rad51* and *rad4 ntg1* mutants exhibit enhanced IQ sensitivities. Synergistic relationship between *NTG1* and *RAD4* has been previously observed for oxidative stress resistance (Gellon *et al*., 2001). Further studies are necessary to determine whether the IQ-N^2^ Gua adduct can be recognized by NER enzymes.

Eleven genes of the 337 IQ resistance genes are associated with colorectal cancer (Table 4). Alleles mapping in human orthologs *RAD18* and *NTHL1*, increase the risk of colorectal cancer. *RAD18 Arg302Gln* increases the colorectal cancer risk in the Japanese patients (Kanzaki *et al*., 2007), while biallelic mutations in *NTHL1*, result in *NTHL1*-associated polyposis (Weren *et al*., 2015). Both *NTG1* and *NTHL1* encode bifunctional glycosylases, containing both glycosylase and lyase activity. *NTG1* is noted to function in a redundant pathway to reduce G:C to T:A transversions, while in humans, formamido-pyrimidine residues, but not 8-oxyG, are recognized by NTHL1 (Das *et al*., 2020). Decreased activity is correlated with a mutation signature 30 (Das *et al*., 2020), where C is converted to T, while increased activity correlates with sequestration of the NER gene *XPG*, the mammalian *RAD2* ortholog. These studies further underscore the primary role of base excision repair and a secondary role of NER in repairing oxidative DNA damage.

In summary, we profiled the yeast genome for IQ resistance using two different yeast libraries; one which was capable of bioactivation and the other which was not. Similarities between the IQ resistance genes indicated that protein was an IQ target, while in the “humanized” library, we identified DNA damage tolerance genes and *NTG1*. Considering that human orthologs *NTHL1* and *RAD18* are risk factors for colon cancer, our results provoke further studies suggesting that polymorphic *NTHL1* alleles may confer HAA sensitivities.

### Data availability

All yeast strains and plasmids are available upon request. Specific yeast strains used in this study are listed in Table S1. Next generation sequencing (NGS) data of barcodes can be found at GEO (https://www.ncbi.nlm.nih.gov/geo/query/acc.cgi?acc=GSE216310). Eleven supplementary tables, five supplementary figures, and a file containing the R code used for processing barcode counts can be accessed on Figshare.

The first file contains the R code for processing barcode counts. Figure S1 shows the activities of CYP1A2 and NAT2 expressed in budding yeast. Figure S2 shows the interaction of proteins encoded by the IQ resistance genes identified in the original yeast deletion collection. Figure S3 shows the competitive growth data for selected diploid strains sensitive to bioactivated IQ. Figure S4 shows the cell viability for selected strains sensitive to bioactivated IQ. Figure S5 shows growth curves of WT and Rad^-^ haploid strains expressing CYP1A2 and NAT2 when exposed to IQ or solvent control.

Table S1 contains the list of specific strains used in this study. Table S2 contains the oligos used for high-throughput sequencing and multiplexing. Table S3 contains mutagenicity data from the Ames assay. Table S4 contains the full list of IQ resistance genes identified from the “humanized” yeast deletion collection in either screen. Table S5 contains the list of IQ resistance genes that were either identified in multiple screens or were independently validated. Table S6 contains the full list of IQ resistance genes identified in the original yeast deletion collection. Table S7 contains the list of IQ resistance genes from the original collection that were identified in multiple screens. Table S8 contains the list of IQ resistance genes which confer resistance with or without bioactivation of IQ. Table S9 contains the full list of genes conferring IQ sensitivity. Table S10 contains the list of GO Process Terms for IQ resistance genes validated or identified in multiple screens with the “humanized” library. Table S11 contains the list of genes conferring resistance to both bioactivated IQ and bioactivated AFB1.

## ACKNOWLEDGMENTS

This work was supported by grant support from the National Institute of Environmental and Health Sciences, R21ES1954 and R15ES023685. We acknowledge Cinzia Cera for assistance in constructing plasmids that express CYP1A2 and NAT2 and Zhenghong Zheng for performing preliminary Western blots using Nat2 antibodies. We thank Chris Vulpe for the original pooled yeast deletion collection library.

## Figure Legends

**Supplemental Figure 1: Measurements of CYP1A2 and NAT2 activities in budding yeast.**

Panel A. CYP1A2 enzymatic activity was measured using the methyl resorufin o-demethylase (MROD) assay. Microsomes were prepared from either wild-type yeast (BY4743) or yeast containing CYP1A2-expression plasmid (BY4743 + pCS316) and MROD activities were measured using rat liver S9 as a positive control and buffer without microsomes as a negative control. X-axis is time in minutes, and y-axis is pmol resorufin released, determined by fluorescence. The table below presents activity readings as pmol of resorufin/mg protein x minute. Panel B. N-acetyltransferase activity of yeast cytosolic protein preps and sulfamethazine (SMZ) with and without acetyl-CoA after 30 minutes incubation at 37°C. Activity is measured as a decrease of SMZ fluorescence. Yeast cytosolic extracts were prepared from the BY4743 strain transformed with either pCYP1A2 or pCYP1A2_NAT2. N=2.

**Supplemental Figure 2: Protein interactome encoded by IQ resistance genes identified in the original “non-humanized” yeast deletion library.**

Panel A. Interactome of the 835 unique ORFs identified from either the IQ resistance screen after 400 μM or 800 μM exposures. Thickness of lines corresponds to evidence of interaction. Singletons and doublets were removed to improve readability. Panel B. Protein interactome of unique ORFs common in both the IQ resistance screen after 400 μM and 800 μM exposures. Nodes with orange fill correspond to the GO Term “Negative regulation of nucleobase-containing compound metabolic process.” Nodes with a blue border correspond to the GO Term “Cellular protein modification process.” Unconnected nodes were removed to improve readability

**Supplemental Figure 3: Competitive growth data for selected diploid strains sensitive to “activated” IQ.** The y-axis shows the percentage of GFP-containing wild type (WT, YB676). Mutant strain and WT were co-inoculated at approximately 10:1 ratio. All strains represented contained pCYP1A2_NAT2. 2×10^4^ cells were acquired per sample. For BY4743, N=8 for MeOH exposure, N =8 for 400 mM IQ exposure, and N=5, for IQ mM 800 exposure. For other strains, N=2 for all treatments. Error bars represent one standard deviation. Significance was determined by either a one-tailed paired Student’s T-Test, or Dunnett’s test, as necessary. Asterisks denote p<0.05 (*), p<0.01(**), and p<0.001 (***).

**Supplemental Figure 4: Cell viability for strains sensitive to activated IQ after treatment with MeOH or IQ.** Cells were treated with either 2% MeOH or 800 mM IQ for 3 hours, then washed three times, resuspended in sterile water, and combined 1:1 with 0.4% trypan blue. Average percent dead cells is determined by dividing the number of cells stained blue by the total number of cells. Total number of cells varied between 200 and 1500. Error bars represent one standard deviation. Significance was determined by either a one-tailed paired Student’s T-Test. Asterisks denote p<0.05 (*), p<0.01(**), and p<0.001 (***).

**Supplemental Figure 5: Growth curves of wild type and Rad^-^ yeast haploid strains expressing CYP1A2 and NAT2 and exposed to IQ.**

Cells were exposed to either MeOH (blue), 400 μM IQ (orange), or 800 μM IQ (gray); growth curves were plotted as absorbance at 600 nm vs time in hours, N=2. Error bars represent one standard deviation. Percent growth for the strains was determined by the ratio of AUCs, AUC_IQ_/AUC_MeOH_. Percent growth for BY4741 was 88% for 400 μM IQ and 84% for 800 μM IQ. Percent growth for *rad4* was 99% for 400 μM IQ and 94% for 800 μM IQ. Percent growth for *rad18* was 43% for 400 μM IQ and 14% for 800 μM IQ. Percent growth for *rad51* was 85% for 400 μM IQ and 77% for 800 μM IQ. Percent growth for *rad4rad51* was 27% for 400 μM IQ and 9% for 800 μM IQ. Percent growth for *rad4rad18* was 64% for 400 μM IQ and 29% for 800 μM IQ. Percent growth for *rad18rad51* was 48% for 400 μM IQ and 23% for 800 μM IQ.

